# ChemEmbed: A deep learning framework for metabolite identification using enhanced MS/MS data and multidimensional molecular embeddings

**DOI:** 10.1101/2025.02.07.637102

**Authors:** Muhammad Faizan-Khan, Roger Giné, Josep M. Badia, Maribel Pérez-Ribera, Alexandra Junza, Maria Vinaixa, Marta Sales-Pardo, Roger Guimerà, Oscar Yanes

## Abstract

Machine learning tools have become essential for annotating the vast number of unidentified MS/MS spectra in metabolomics, addressing the limitations of current reference spectral libraries. However, these tools often struggle with the high dimensionality and sparsity of MS/MS spectra and metabolite structures. ChemEmbed introduces a novel approach by combining multidimensional and continuous vector representations of chemical structures with enhanced MS/MS spectra. This enhancement is achieved by merging spectra from multiple collision energies and incorporating calculated neutral losses from 38,472 distinct compounds, providing richer input for a convolutional neural network (CNN). ChemEmbed achieves top-ranked candidate annotations in over 42% of cases and identifies the correct compound within the top five in more than 76% of cases in a test dataset. Against external benchmarks such as CASMI 2016 and 2022, ChemEmbed outperforms SIRIUS, the current state-of-the-art in computational metabolomics. In a validation experiment with the Annotated Recurrent Unidentified Spectra (ARUS) dataset— including over 25,000 spectra from human plasma and 68,000 from urine— ChemEmbed successfully identified 24 previously unannotated compounds. By aligning with the advanced capabilities of modern mass spectrometry instrumentation, ChemEmbed balances accuracy, computational efficiency, and scalability, making it a powerful solution for high-throughput metabolomics applications.

## Introduction

The interpretation of tandem mass spectrometry (MS/MS) data is crucial for the identification of metabolites, which in turn is particularly relevant for the growing applications of metabolomics in biomedicine, nutritional and environmental sciences^1^.

The predominant strategy for metabolite identification involves comparing experimental spectra with pre-recorded MS/MS spectra of known compounds to find matching fragments. However, due to the limited size, quality, and diversity of available reference spectral libraries, a significant portion of MS/MS spectra generated in metabolomic experiments remain unidentified^2^.

*In silico* fragmentation tools have emerged as a solution to address this challenge. These tools combine chemical rules to capture known fragmentation events^3^, probabilistic models to assign probabilities to potential fragmentations^3,4^, and/or machine learning algorithms to learn the complex relationships between molecular structures and their corresponding fragmentation patterns by MS^5^.

To enable efficient computation and learning by machine learning algorithms, both experimental MS/MS spectra and the chemical structures of metabolites must be converted into numerical formats that preserve key information. MS/MS spectra, typically represented as lists of m/z values and intensities, are encoded into fixed-length vectors using techniques such as feature hashing^6,7^, binning^8–10^, and spectral fingerprinting^11^. Meanwhile, chemical structures are typically encoded as molecular fingerprints^12–14^, though graph-based representations are also used^15,16^.

Despite these encoding techniques^16^, spectral and structural data in metabolomics still presents challenges due to its high dimensionality and sparsity. These issues contribute to problems like model interpretability, computational complexity, and overfitting, making it difficult for machine learning algorithms to learn meaningful patterns and generalize effectively. Molecular fingerprints, for example, are binary or count vectors that represent the presence, absence, or the number of specific chemical features within a molecule, such as atom types, functional groups, or structural motifs. While the dimensionality of these fingerprints is fixed by the number of features they are designed to capture, the vast chemical diversity of metabolites leads to inherent sparsity—most features remain inactive or set to zero for most compounds. Similarly, encoded m/z values and intensities from MS/MS spectra face the same sparsity issue when represented as fixed-length vectors. Given that spectra only contain a few peaks within a broad m/z range, most bins are left empty, with only a small number of non-zero bins corresponding to specific fragment ions.

To overcome these limitations, we propose ChemEmbed, a new tool for metabolite identification from MS/MS spectra. Our tool is based on two complementary strategies designed to enrich the representations of both chemical structures and MS/MS spectra. First, we employ 300-dimensional embeddings generated by Mol2vec^17^, an unsupervised machine learning approach that captures complex structural properties of molecules with greater depth than traditional molecular fingerprints. Second, to address the sparsity in spectral data, we use merged spectra that combine multiple collision energies and calculated neutral losses from reference libraries. This expanded spectral representation broadens the range and diversity of binned spectra, providing a richer input for a convolutional neural network (CNN). The CNN is trained to predict 300-dimensional Mol2vec embeddings from these enhanced spectra, and these embeddings are then compared and ranked against a reference database of millions of Mol2vec embeddings.

We consider a dataset of 38,472 distinct compounds with reference MS/MS data, which we split into training, validation, and test sets. In the test set, ChemEmbed accurately annotated the top-ranked candidate (Tanimoto ≥ 0.95) in over 42% of cases and identified the correct candidate within the top five in more than 76% of cases. When challenged with external datasets such as CASMI 2016 and 2022, ChemEmbed consistently outperformed the latest version of SIRIUS^11^, the current state-of-the-art method in computational metabolomics. Furthermore, in a validation experiment using the Annotated Recurrent Unidentified Spectra (ARUS) from the NIST Mass Spectrometry Data Center, our tool successfully identified 24 previously unannotated compounds, demonstrating its broad applicability and superior performance across diverse datasets.

## Results

### Construction of spectral and structural databases

We generated SMILES identifiers for 38,472 unique compounds sourced from the NIST20^18^, MSDIAL^19^, GNPS^20^, and Agilent METLIN metabolomics libraries (see Methods) (Supplementary File 1), comprising 683,664 MS/MS spectra in positive ionization ([M+H]^+^) and 158,382 MS/MS spectra in negative ionization ([M-H]^−^), as detailed in Supplementary Tables 1 and 2, respectively. Since many compounds were annotated across different reference libraries, we calculated the overlap by matching the first 14 characters of their InChIKey identifiers (Supplementary Figure 1). To resolve duplicates, we prioritized the libraries in the following order: NIST20 > Agilent METLIN > GNPS > MSDIAL (Supplementary Table 3 and 4). For example, if a compound was found in both NIST20 and Agilent METLIN, we discarded the MS/MS spectra from Agilent METLIN, and so on.

These spectra were pre-processed and curated to remove background signals and noise, as described in the Methods section. This collection formed what we termed the individual spectra dataset, capturing the diversity and redundancy of compounds fragmented at multiple collision energies.

Next, we created a second collection of MS/MS spectra by merging all available spectra across different collision energies for each compound, producing a single representative MS/MS spectrum per compound. This merging process resulted in 38,472 spectra in positive ionization mode and 14,168 spectra in negative ionization mode. The merged spectra dataset broadened the range and diversity of encoded bins, providing a more comprehensive view of each compound’s fragmentation behavior by combining information from various collision energies.

To further enhance spectral representation, we generated a third collection of MS/MS spectra by incorporating neutral losses (NL) into the merged spectra. For both positive and negative ionization modes, we calculated the mass difference between precursor and fragment ions, and then binarized their intensities, rather than retaining the original fragment ion intensities, as done in previous studies^21^. The resulting neutral loss values were added to each merged spectrum, resulting in an enriched dataset that combines fragment ions from multiple collision energies with their corresponding neutral losses, all in binarized format.

### Mol2vec embeddings reference database

Mol2vec is an unsupervised deep learning model, trained on 19.9 million compounds from the ZINC^22^ and ChEMBL^23^ databases, which converts molecular structures from their SMILES into 300-dimensional numerical vector embeddings. Since a single InChIKey can produce multiple redundant SMILES, and our goal was to use Mol2vec embeddings as labels for training the CNN model, we needed to confirm that Mol2vec was insensitive to different SMILES representations. To verify this, we randomly selected 100 compounds from our database, generated 10 compatible SMILES for each, and compared the resulting 300-dimensional embeddings. The Euclidean distance between the embeddings was nearly zero (data not shown), confirming that Mol2vec embeddings are consistent across SMILES versions and can serve as reliable labels for CNN model training.

Following this validation, we generated a comprehensive reference database of 0.52 million molecules, each with its corresponding Mol2vec embedding, sourced from COCONUT^24^, HMDB^25^, NIST20, GNPS, Agilent METLIN, and MSDial. These embeddings serve as the “ground truth” or reference embeddings for subsequent comparisons in the CNN model (Supplementary File 2).

### CNN model for molecular embedding prediction from MS/MS data

We developed a convolutional neural network (CNN) (Supplementary Figure 2) that accepts MS/MS spectra as input and generates 300-dimensional embeddings as output. The CNN’s primary objective is to map the input spectra into a molecular embedding space that replicates the Mol2vec representations. For each spectrum in the test dataset, the predicted embeddings were then compared to reference database of 0.52 million molecules, each represented by its Mol2vec embedding. To ensure accurate matching between the predicted embeddings and known molecules, we employed a multi-step filtering and ranking strategy (Figure 1).

**Figure 1.**
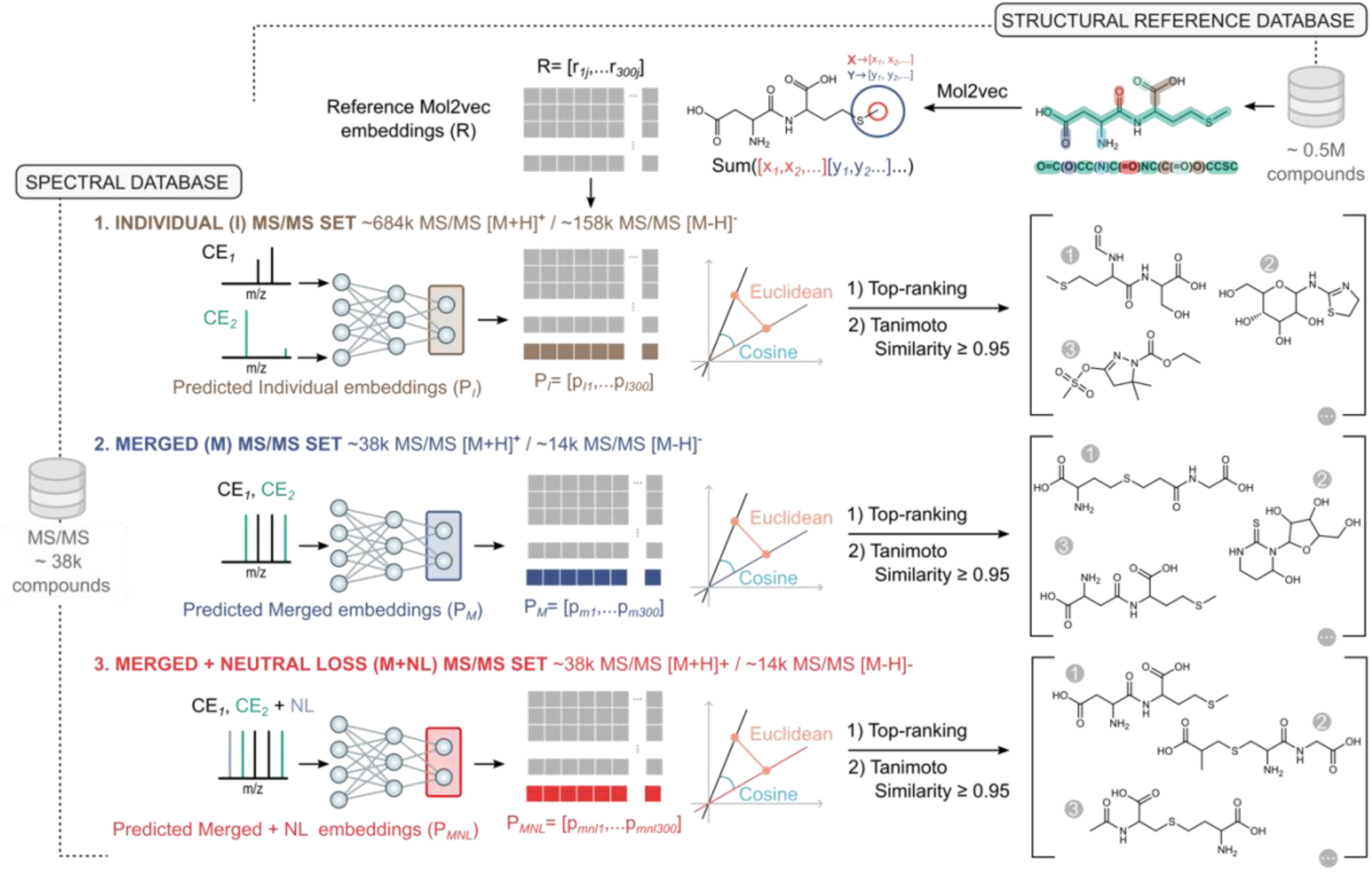
Computational workflow for molecular identification using CNN-generated embeddings. MS/MS spectra are the input to a convolutional neural network (CNN) trained to generate 300-dimensional embeddings aligned with Mol2vec representations. Predicted embeddings are compared to a reference database in a multi-step process: (i) Precursor mass filtering: Candidate molecules with precursor ion masses outside a ±0.001 Da error tolerance are excluded; (ii) Similarity calculations: Euclidean distance and cosine similarity are computed between the predicted embeddings and reference Mol2vec embeddings, ranking molecules by similarity; (iii) Structural validation: The top-ranked candidates are further evaluated using the Tanimoto score, ensuring structural alignment between predicted and reference molecules. A Tanimoto score threshold of ≥0.95 is applied to confirm molecular identity.

First, we applied a precursor ion mass filter to exclude any precursor ions outside a mass error tolerance of ±0.001, focusing the comparison on candidate molecules with closely matching mass. Following this, we calculated both the Euclidean distance and cosine similarity between the predicted 300-dimensional vector representation and the 0.52 million reference Mol2vec embeddings. Molecules were then ranked by their similarity, with Euclidean distance providing a measure of closeness between two points in the embedding space, and cosine similarity capturing the directional alignment between the predicted and reference vectors. Together, they provide a more comprehensive assessment of similarity, capturing both spatial proximity and pattern alignment, enhancing the reliability of the molecular ranking process.

Finally, to assess the structural similarity between the predicted and reference molecules, we calculated the Tanimoto score for the top-ranked molecules based on their Euclidean distance and cosine similarity. The Tanimoto score ranges from 0 to 1, where 0 indicates no similarity and 1 indicates perfect similarity. In cheminformatics, thresholds above 0.85 are often regarded as indicative of significant structural similarity^26–28^. Here, we set a threshold of 0.95 for the Tanimoto score to determine molecular identity. This threshold provided an additional layer of validation, ensuring that the predicted embedding not only closely matched the reference embedding but also corresponded to a structurally very similar (e.g., stereoisomers) or identical molecule in terms of their SMILES representations.

### Performance of ChemEmbed in a test dataset

The three collections of MS/MS spectra—individual, merged, and merged with neutral losses—were used to train and test three CNN models to predict molecular embeddings, with 80% of the spectral data used for training, 10% for validation, and 10% for testing (see Methods).

Figure 2A displays the distributions of Euclidean distances and cosine similarities between the predicted 300-dimensional embeddings and the reference Mol2vec embeddings for the individual spectra dataset in positive ionization. The model trained on this dataset showed a mean Euclidean distance of 40 and a mean cosine similarity of 0.93. In comparison, the model trained on the merged spectra dataset without neutral losses exhibited a significantly lower mean Euclidean distance of 24.5 and a higher mean cosine similarity of 0.96. Incorporating neutral losses provided only a slight improvement, reducing the Euclidean distance to 23.1 and increasing the cosine similarity to 0.97. These findings demonstrate that as the richness of spectral information increases—by merging spectra and including neutral losses—the CNN’s predicted embeddings more closely align with the ground truth Mol2vec embeddings. Similar improvements were observed in the negative ionization mode (Figure 2B).

**Figure 2.**
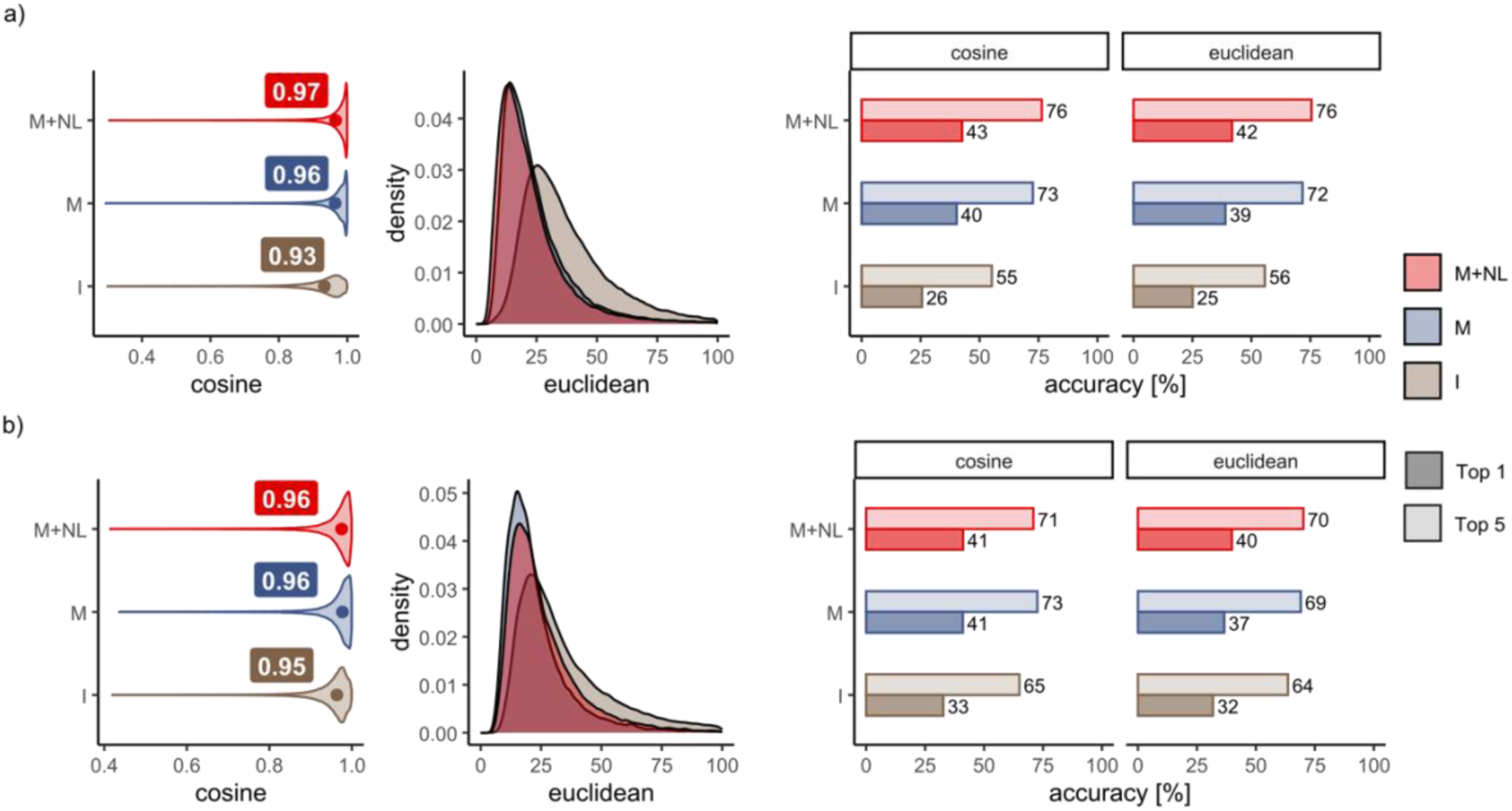
Alignment of CNN-predicted embeddings with reference Mol2vec embeddings. Violin and density plots illustrate the distributions of cosine similarities and Euclidean distances between the CNN-predicted 300-dimensional embeddings and the reference Mol2vec embeddings. Results are shown for positive ionization (**a**) and negative ionization (**b**) datasets. Three model configurations are compared: trained with individual spectra (I), trained with merged spectra (M), and trained with merged spectra incorporating neutral losses (M+NL). Accuracy metrics are represented as bars, indicating the percentage of correctly identified molecules ranked in the top 1 (dark color) and top 5 (light color) positions for both ionization modes. The plots highlight the performance improvements associated with merging spectra and incorporating neutral losses.

This improvement in embedding accuracy directly impacted the model’s ability to rank correct compound annotations. When trained on the individual spectra dataset, the CNN correctly annotated the top-ranked candidate (Tanimoto score ≥0.95) in over 26% of cases and identified the correct compound within the top five ranked candidates in over 56% of cases. In contrast, the model trained on the merged spectra dataset significantly improved annotation performance, with the correct top-ranked candidate identified in over 40% of cases and the correct compound found within the top five in over 73% of cases. The inclusion of neutral losses further enhanced CNN’s performance, with correct top-ranked candidates identified in over 43% of cases and correct annotations within the top five in over 76% of cases.

Performance in the negative ionization mode showed consistent and reproducible behavior, with only a slightly slower ranking performance compared to positive ionization (Figure 2B). We attribute this difference to the typically lower number of fragment ions in negative ionization, which may provide less informative data for the CNN to generate accurate molecular embeddings.

As expected, neither the combination of individual mass spectra with neutral losses (59.8% for top 5 annotations) nor neutral losses alone (69% for top 5 annotations, calculated from merged spectra) outperformed the full combination of merged spectra with neutral losses (Supplementary Table 5).

An additional advantage of using merged spectra is that the processing time for the test dataset, which included 3,850 compounds, was substantially reduced with respect to considering individual spectra. On a basic laptop CPU, the full pipeline—covering MS/MS preprocessing, CNN inference, and candidate ranking—took less than 20 minutes for the merged spectra with neutral losses. In contrast, processing the individual spectra dataset required nearly 6 hours, highlighting the efficiency and scalability of the merged spectra approach.

To further highlight the importance of combining enhanced MS/MS data with multidimensional molecular embeddings, we investigated molecular fingerprints as an alternative encoding method for chemical structures. Specifically, we employed molecular fingerprints encompassing 4,237 molecular properties (see Methods), like those used in SIRIUS-CSI:FingerID^29^, within our CNN architecture. Replacing Mol2vec embeddings with fingerprints failed to show significant learning. Instead, fingerprints performed better with a deep neural network (DNN) architecture comprising four fully connected layers. This approach, however, resulted in lower performance, with correct top-ranked candidates identified in 39% of cases and correct annotations within the top five in 73% of cases (Supplementary Table 6).

### Validation with non-annotated spectra

To evaluate the tool with datasets of non-annotated metabolites, we used three publicly available resources: the Critical Assessment of Small Molecule Identification (CASMI) challenges^30^ from 2016 and 2022, as well as the Annotated Recurrent Unidentified Spectra (ARUS) database^31^ from the NIST Mass Spectrometry Data Center. To expand the pool of potential candidate structures, we created a reference database containing Mol2vec embeddings for 5.52 million molecules, which included 5 million random compounds from PubChem.

The CASMI 2016 challenge dataset contained 208 MS/MS spectra from 188 unique structures, with 127 spectra acquired in positive ion mode and 81 in negative ion mode. After removing any molecules from CASMI 2016 that were present in our CNN training dataset, we retained 27 unique structures in positive mode and 30 in negative mode. For CASMI 2022, which initially included 177 structures in positive ion mode and 108 in negative ion mode, we applied the same filtering process. This resulted in 149 unique compounds in positive mode and 97 in negative mode.

The ARUS dataset contains MS/MS spectra frequently observed in human samples, but which remain unannotated. We selected ARUS spectra with putative molecular formulas assigned using the BUDDY software^32^ (see Supplementary File 3). This dataset includes 25,801 spectra from plasma and 68,478 spectra from urine.

For each unknown molecule, we pre-processed its associated spectrum as previously described (see Methods). That is, peaks exceeding the precursor mass by more than 0.5 Da were removed, and fragments with intensities below 1% of the most intense peak were filtered out. The mass differences between precursor and fragment ions were calculated, and the resulting neutral loss values were incorporated into each spectrum. All intensity values were binarized and each spectrum was then encoded as a vector with a bin size of 0.01 Da, with the m/z axis restricted to ≤ 700 Da.

With our tool, we generated a list of candidate annotations for each spectrum. These candidates were ranked based on both the Euclidean distance and cosine similarity between the predicted 300-dimensional embeddings and a reference set of 5.52 million Mol2vec embeddings. The top five candidates were selected for further analysis.

In the CASMI 2016 challenge, our model successfully ranked the correct molecule within the top five candidates in over 60% of cases, for both positive and negative ionization spectra (Supplementary Table 7).

We then compared the performance of ChemEmbed against the latest version of SIRIUS (version 6.0.0)^11^ (see Methods) using both the CASMI 2022 and ARUS datasets. Both tools used the same reference dataset of 5.52 million molecules as the pool of potential candidate structures, matching either molecular fingerprints (SIRIUS) or 300-dimensional Mol2vec embeddings ChemEmbed. To ensure a fair comparison, we removed compounds from the CASMI 2022 dataset that had been included in the training of the latest version of SIRIUS (see Methods), resulting in 107 unknown compounds for both tools in positive ionization mode and 64 in negative ionization mode. ChemEmbed ranked the correct compound within the top five candidates in 33% of cases for positive ionization spectra and 30% for negative ionization spectra. In contrast, SIRIUS achieved success rates of 28% for positive ionization and 17% for negative ionization when considering the top five candidates (Supplementary Table 7).

Finally, we applied ChemEmbed to process 25,801 MS/MS spectra from plasma and 68,478 spectra from urine in the ARUS dataset, generating potential annotations for 23.8% of positive ionization spectra (Supplementary File 4) and 19.7% of negative ionization spectra (Supplementary File 5) with cosine similarity scores above 0.95 and Euclidean distances below 25 (Supplementary Table 8). From these, we randomly selected 40 spectra—10 from each biological matrix and ionization mode. An analytical chemistry expert then manually reviewed the spectra to identify which of the top five candidates best matched the fragmentation patterns, leading to the identification of 24 compounds: 18 in positive ionization mode and 6 in negative.

Attempts to replicate this analysis using SIRIUS 6.0 were unsuccessful, as the tool either failed to complete the processing of the large dataset or caused the server to crash. As a result, we limited the comparison to the 18 compounds identified in positive ionization mode across plasma and urine. Of the 18 compounds annotated by ChemEmbed, SIRIUS correctly identified 14 within the top five. However, only three of these were ranked higher by SIRIUS compared to ChemEmbed (Figure 3).

**Figure 3.**
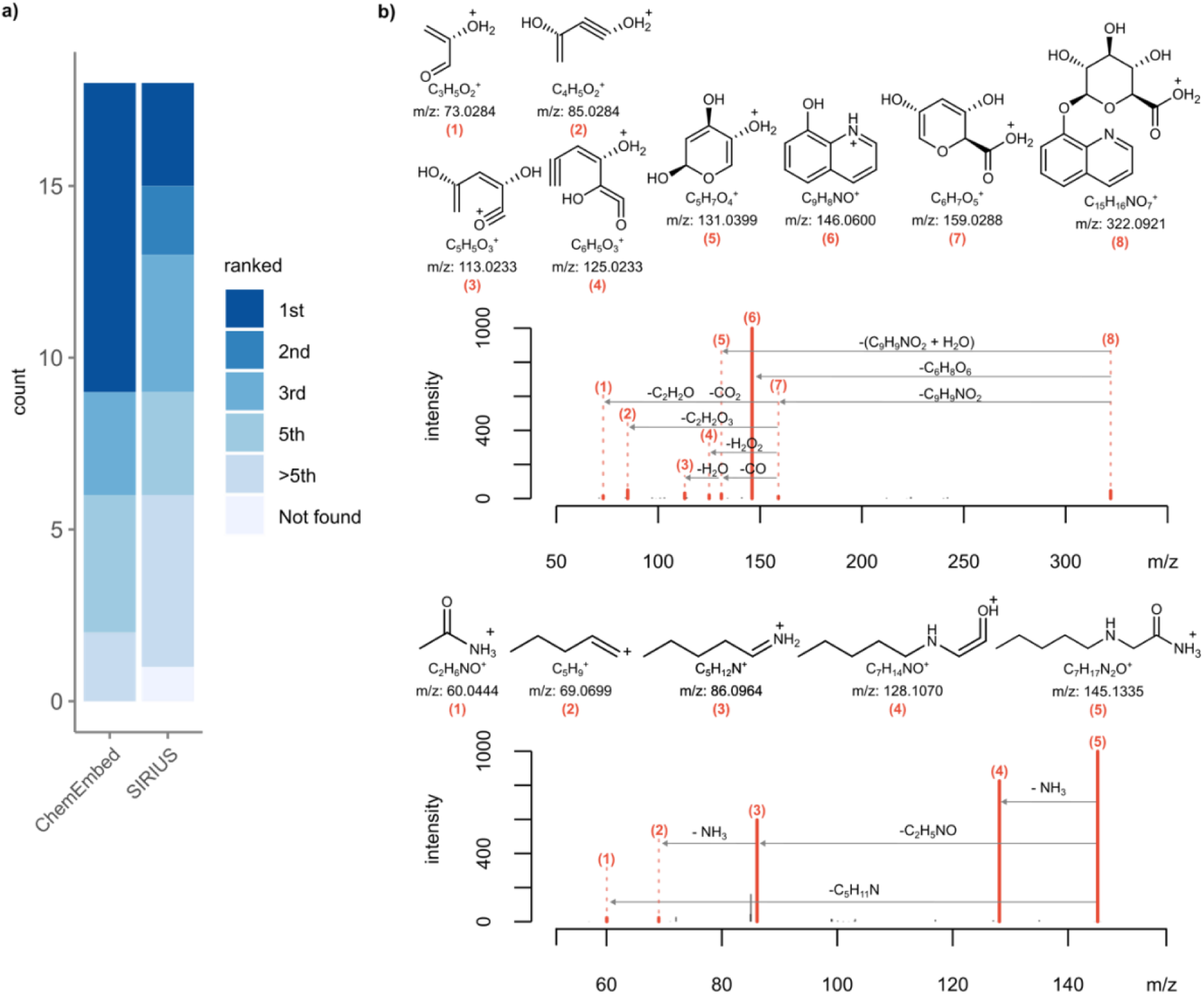
ChemEmbed applied to the ARUS dataset. (**a**) Stacked bar chart showing the ranking positions of 18 compounds identified in positive ionization mode across plasma and urine by ChemEmbed, compared to SIRIUS. ChemEmbed correctly identified all 18 compounds within the top five rankings, whereas SIRIUS identified 14. Notably, SIRIUS ranked only three compounds higher than ChemEmbed. (**b**) MS/MS spectra elucidation for 8-hydroxyquinoline glucuronide (top) and Milacemide (bottom) from the ARUS urine spectral dataset, displaying tentative fragment ion annotations and molecular losses from precursor to product ions.

## Discussion

Despite the growing number of reference MS/MS spectra, public and commercial databases still cover less than 5% of the known small molecule chemical space^33,34^. Even focusing on genome-scale metabolic networks—excluding the vast chemical diversity introduced by microbiota, environmental exposures, diet, and contaminants—only about 40% of eukaryotic metabolic networks can be mapped using available spectral standards^35^.

Machine learning tools have been developed to bridge this gap, but they face challenges in learning meaningful patterns due to the high dimensionality and sparsity of MS/MS spectra and metabolite structures. A key issue stems from training models on reference MS/MS spectra acquired at fixed collision energies, as seen in foundational databases like METLIN, MassBank, and GNPS. Our study demonstrates that training neural networks with individual collision energy spectra per molecule underperforms compared to using a single merged spectrum. The higher sparsity of individual spectra complicates learning and increases the risk of overfitting.

Using merged spectra also addresses the increased dimensionality and computational complexity of fixed-energy inputs. For instance, training one epoch with individual spectra took over 95 minutes, compared to just 5.5 minutes with merged spectra. This efficiency extends to inference: processing 25,801 MS/MS spectra from plasma and 68,478 from urine in the ARUS dataset required less than 1 hour and 2 hours on a basic laptop CPU, respectively. These results highlight the scalability and practicality of our approach.

Modern qTOF and Orbitrap instruments now support high scan rates, improved duty cycles, and advanced collision energy options, such as full-CE ramps and stepped collision energies. These advancements provide more comprehensive fragmentation patterns, enhancing metabolite identification accuracy. However, leveraging such rich spectral information requires careful preprocessing. Our tool, ChemEmbed, merges data from multiple collision energies and binarizes the intensities, simplifying the input and reducing overfitting risks by focusing on the presence of fragment ions rather than their intensities. Fragment intensities can vary widely due to factors such as fragmentation technique, collision energy, and instrument-specific differences^36,37^ adding unnecessary complexity and potential noise if treated as model inputs.

Our results also emphasize the importance of aligning neural network architectures with the nature of molecular representations. CNNs paired with Mol2vec embeddings excel in capturing the complex relationships within high-dimensional, continuous vector representations. In contrast, the sparse, binary nature of molecular fingerprints limits their utility in CNNs, favoring simpler architectures like DNNs.

Additionally, while the limited availability of reference spectra remains a primary bottleneck^38^, the quality of MS/MS data in public (e.i., community-contributed) and commercial libraries is equally critical. High-quality datasets, such as the NIST library, set a benchmark that others should follow to improve computational tools’ performance.

In summary, ChemEmbed aligns with the capabilities of modern mass spectrometry instrumentation by balancing performance metrics like accuracy, computational cost, and scalability. Unlike many prior tools, it addresses real-world usability, making it a robust solution for metabolite identification in high-throughput applications.

## Methods

### 1. Dataset and data preprocessing

For our spectra-to-molecule convolutional neural network (CNN) model, we used a dataset of 38,472 unique compounds sourced from the NIST20, MSDIAL, GNPS, and Agilent METLIN databases. Initially, the NIST20 dataset provided only InChIKey information, so we leveraged the PubChem API to retrieve the corresponding SMILES strings. The Agilent METLIN dataset also contained missing SMILES and InChIKey data, which we supplemented using the RDKit tool to convert available SMILES to molecular information, followed by PubChem API queries to obtain the remaining SMILES for specific InChIKeys.

After gathering the SMILES information for all datasets, we removed any spectra with null values. Since stereochemistry was not considered in this work, we used only the first 14 characters of the unique InChIKey identifier.

### 2. Spectral data preprocessing

To prepare the MS/MS spectra for model training, several preprocessing steps were implemented. Only [M+H]^+^ adducts in positive ionization and [M-H]^−^ adducts in negative ionization were used. Peaks that exceeded the precursor mass by more than 0.5 Da were removed. To reduce noise, fragments with intensities below 1% of the highest peak were filtered out, and the remaining intensities were binarized. Each MS/MS spectrum was encoded as a vector with a bin size of 0.01. Pareto analysis revealed that 20% of the unique m/z bins accounted for 83% of the spectral values (Supplementary Figure 3). Consequently, we limited the m/z axis to ≤ 700, reducing the vector length to 70,000 bins. This adjustment eased computational demands while retaining 80% of the spectral data.

### 3. Convolutional Neural Networks (CNNs)

We employed a convolutional neural network (CNN) to predict 300-dimensional Mol2vec embeddings from input MS/MS spectra. The models were trained and tested using an NVIDIA Tesla T4 GPU, with experiments conducted within the PyTorch framework. For development and implementation, we utilized the PyCharm integrated development environment (IDE).

The dataset of individual spectra was split into three subsets: 527,236 spectra (80%) for training, 64,774 spectra (10%) for validation, and 66,843 spectra (10%) for testing. Similarly, the merged spectra dataset was divided into 30,775 spectra (80%) for training, 3,847 spectra (10%) for validation, and 3,850 spectra (10%) for testing.

#### 3.1 Network architecture

Six convolutional layers capture spatial relationships between mass-to-charge (m/z) values across the spectrum. Six max pooling layers reduce the dimensionality of the feature space. The first fully connected layer flattens the spatial data, and the final fully connected layer outputs the 300-dimensional Mol2vec embedding. The input to the network is a vector of 70,000 bins, representing the m/z values of the spectra.

#### 3.2. Hyperparameters and training

We used the Adam optimizer with a learning rate of 0.0001 for backpropagation. Early stopping based on validation loss was implemented to prevent overfitting. The merged spectra network was trained for 70 epochs, while the individual spectra network was trained for 15 epochs, both using a mini-batch size of 32. Mean squared error was used as the cost function.

#### 3.3. Activation function and regularization

The Rectified Linear Unit (ReLU) was chosen as the activation function due to its effectiveness in mitigating the vanishing gradient problem in deep networks. Dropout regularization was applied to prevent overfitting. No activation function was applied to the final output layer, allowing the predicted embeddings to span any value within the Mol2vec embedding space.

### 4. Analysis with SIRIUS 6.0

To validate our tool against SIRIUS+CSI FingerID, we used the graphical interface of SIRIUS v6.0.0 (June 3, 2024). First, we imported a custom reference database of 5.52 million entries into the software using SMILES information to ensure consistency across both tools. Next, we removed chemical structures from CASMI 2022 that were part of the SIRIUS 6.0 training set by downloading the InChI identifiers for positive and negative ion mode from the respective training structure links: https://csi.bright-giant.com/v3.0/api/fingerid/trainingstructures?predictor=1 and https://csi.bright-giant.com/v3.0/api/fingerid/trainingstructures?predictor=2. We then converted these InChI identifiers to SMILES format using RDKit.

We imported the MS/MS spectra from the CASMI 2022 dataset and initiated the analysis by selecting the “Compute” option, which opened a parameter configuration window. For the SIRIUS tool, we configured the instrument as ORBITRAP, MS2 mass accuracy to 5 ppm, the adduct to [M+H]^+^ and [M-H]^−^ for positive and negative ion mode, respectively, and set the molecular formula generation method to De novo + bottom-up. Following this, we enabled CSI:FingerID for property prediction with default threshold score settings. For the structure database search, we selected our custom database of 5.52 million structures and disabled PubChem as a fallback. Upon running the analysis, SIRIUS returned a list of candidate molecules for each MS/MS spectrum, which we sorted in ascending order by the CSI:FingerID score, a key feature used to assess the quality of candidate molecules.

In the final step, we identified the top 5 candidates from both tools and compared their performance by evaluating which tool returned a higher number of correct candidates in the top 5 for each MS/MS spectrum. This comparison enabled us to assess the relative accuracy and effectiveness of the two tools in generating valid candidate molecules.

### 5. Molecular Fingerprints

Molecular fingerprints, commonly used to represent molecular structures, encode predefined molecular features, such as specific substructures, into bit vectors. To replace Mol2vec embeddings, we used all five fingerprint types defined in the SIRIUS CSI framework^39^: (i) CDK Substructure Fingerprints: Capture 307 molecular properties using predefined substructure patterns (Chemistry Development Kit, version 1.5.8); (ii) PubChem (CACTVS) Fingerprints: Represent 881 properties, based on PubChem specifications; (iii) Klekota–Roth Fingerprints: cover 4,860 properties, providing detailed structural and functional group information; (iv) FP3 Fingerprints: represent 55 properties derived from SMARTS patterns (Open Babel, version 2.3.2); and (v) MACCS Fingerprints: Encode 166 SMARTS-based properties (Open Babel, version 2.3.2). Fingerprint calculations utilized the Chemistry Development Kit (CDK) version 1.5.8 and PyFingerprint (https://pypi.org/project/pyfingerprint/).

#### 5.1 Data preprocessing

The initial combined fingerprint set encompassed 6,269 molecular properties. Constant and redundant features were removed from the training dataset, reducing the feature set to 4,237 properties. These features were split into training (80%), validation (10%), and testing (10%) subsets, maintaining the same distribution used for Mol2vec-based training.

#### 5.2 Model development and training

Initial attempts to train the CNN architecture designed for Mol2vec embeddings with molecular fingerprints failed, as training and validation losses remained constant. To address this, a deep neural network (DNN) comprising four fully connected layers with ReLU activation functions was implemented. Dropout layers were added for regularization, and binary cross-entropy served as the loss function to handle the multi-label classification task. During training, outputs were binarized using a threshold of 0.50, assigning a label of 1 to predictions ≥0.50 and 0 otherwise. The DNN demonstrated learning, with steadily decreasing training and validation losses.

#### 5.3 Molecular ranking and evaluation

The trained model was evaluated using the same molecular ranking method as for Mol2vec-based predictions. A dataset of 0.52 million molecules was used, substituting the refined molecular fingerprints for Mol2vec embeddings. Model performance was assessed by calculating cosine similarity between predicted and ground truth fingerprints.

## Data availability

All Supplementary Files are available on the Zenodo repository at: https://zenodo.org/records/14778996.

## Code availability

A Python implementation of ChemEmbed and trained ChemEmbed models are available from https://github.com/massspecdl/ChemEmbed

## Author contributions statement

MS-P, RGu and OY designed the research. MFK performed experiments and wrote code. MFK, MP-R, RGi and JMB collected and processed spectral data. SJ performed the manual elucidation of ARUS spectra. MFK, MS-P, RGu and OY analyzed and discussed results. MFK and OY wrote the paper. MFK and MV designed the figures. All authors commented on the manuscript.

## References

1. Giera, M., Yanes, O. & Siuzdak, G. Metabolite discovery: Biochemistry’s scientific driver. Cell Metab 34, 21–34 (2022).

2. Wishart, D. S. et al. HMDB 5.0: the Human Metabolome Database for 2022. Nucleic Acids Res 50, D622–D631 (2022).

3. Ruttkies, C., Neumann, S. & Posch, S. Improving MetFrag with statistical learning of fragment annotations. BMC Bioinformatics 20, 1–14 (2019).

4. Tsugawa, H. et al. Hydrogen Rearrangement Rules: Computational MS/MS Fragmentation and Structure Elucidation Using MS-FINDER Software. Anal Chem 88, 7946–7958 (2016).

5. Dührkop, K. et al. SIRIUS 4: a rapid tool for turning tandem mass spectra into metabolite structure information. Nat Methods 16, 299–302 (2019).

6. Wolf, S., Schmidt, S., Müller-Hannemann, M. & Neumann, S. In silico fragmentation for computer assisted identification of metabolite mass spectra. BMC Bioinformatics 11, 1–12 (2010).

7. Bittremieux, W., Laukens, K. & Noble, W. S. Extremely Fast and Accurate Open Modification Spectral Library Searching of High-Resolution Mass Spectra Using Feature Hashing and Graphics Processing Units. J Proteome Res 18, 3792–3799 (2019).

8. Van Der Hooft, J. J. J., Wandy, J., Barrett, M. P., Burgess, K. E. V. & Rogers, S. Topic modeling for untargeted substructure exploration in metabolomics. Proc Natl Acad Sci U S A 113, 13738–13743 (2016).

9. Fan, Z., Alley, A., Ghaffari, K. & Ressom, H. W. MetFID: artificial neural network-based compound fingerprint prediction for metabolite annotation. Metabolomics 16, (2020).

10. Smith, C. A., Want, E. J., O’Maille, G., Abagyan, R. & Siuzdak, G. XCMS: Processing mass spectrometry data for metabolite profiling using nonlinear peak alignment, matching, and identification. Anal Chem 78, 779–787 (2006).

11. Dührkop, K. et al. SIRIUS 4: a rapid tool for turning tandem mass spectra into metabolite structure information. Nature Methods 2019 16:4 16, 299–302 (2019).

12. Baygi, S. F. & Barupal, D. K. IDSL_MINT: a deep learning framework to predict molecular fingerprints from mass spectra. J Cheminform 16, 1–8 (2024).

13. Laponogov, I., Sadawi, N., Galea, D., Mirnezami, R. & Veselkov, K. A. ChemDistiller: an engine for metabolite annotation in mass spectrometry. Bioinformatics 34, 2096–2102 (2018).

14. Dührkop, K. Deep kernel learning improves molecular fingerprint prediction from tandem mass spectra. Bioinformatics 38, i342 (2022).

15. Li, S. et al. Encoding the atomic structure for machine learning in materials science. Wiley Interdiscip Rev Comput Mol Sci 12, e1558 (2022).

16. Huber, F., van der Burg, S., van der Hooft, J. J. J. & Ridder, L. MS2DeepScore: a novel deep learning similarity measure to compare tandem mass spectra. J Cheminform 13, 1–14 (2021).

17. Jaeger, S., Fulle, S. & Turk, S. Mol2vec: Unsupervised Machine Learning Approach with Chemical Intuition. J Chem Inf Model 58, 27–35 (2018).

18. Wallace, W. E. & Moorthy, A. S. NIST Mass Spectrometry Data Center Standard Reference Libraries and Software Tools: Application to Seized Drug Analysis. J Forensic Sci 68, 1484 (2023).

19. Tsugawa, H. et al. MS-DIAL: data-independent MS/MS deconvolution for comprehensive metabolome analysis. Nature Methods 2015 12:6 12, 523–526 (2015).

20. Wang, M. et al. Sharing and community curation of mass spectrometry data with Global Natural Products Social Molecular Networking. Nature Biotechnology 2016 34:8 34, 828–837 (2016).

21. Aisporna, A. et al. Neutral Loss Mass Spectral Data Enhances Molecular Similarity Analysis in METLIN. J Am Soc Mass Spectrom 33, 530–534 (2022).

22. Irwin, J. J. et al. ZINC20 - A Free Ultralarge-Scale Chemical Database for Ligand Discovery. J Chem Inf Model 60, 6065–6073 (2020).

23. Zdrazil, B. et al. The ChEMBL Database in 2023: a drug discovery platform spanning multiple bioactivity data types and time periods. Nucleic Acids Res 52, D1180–D1192 (2024).

24. Sorokina, M., Merseburger, P., Rajan, K., Yirik, M. A. & Steinbeck, C. COCONUT online: Collection of Open Natural Products database. J Cheminform 13, 1–13 (2021).

25. Wishart, D. S. et al. HMDB 5.0: the Human Metabolome Database for 2022. Nucleic Acids Res 50, D622–D631 (2022).

26. Zulfiqar, M. et al. Untargeted metabolomics to expand the chemical space of the marine diatom Skeletonema marinoi. Front Microbiol 14, (2023).

27. Dunkel, M. SuperNatural: a searchable database of available natural compounds. Nucleic Acids Res 34, D678–D683 (2006).

28. Nuzzo, A. et al. Expanding the drug discovery space with predicted metabolite– target interactions. Commun Biol 4, 288 (2021).

29. Dührkop, K., Shen, H., Meusel, M., Rousu, J. & Böcker, S. Searching molecular structure databases with tandem mass spectra using CSI:FingerID. Proc Natl Acad Sci U S A 112, 12580–12585 (2015).

30. McEachran, A. D. et al. Revisiting Five Years of CASMI Contests with EPA Identification Tools. Metabolites 10, 260 (2020).

31. Simón-Manso, Y. et al. Mass Spectrometry Fingerprints of Small-Molecule Metabolites in Biofluids: Building a Spectral Library of Recurrent Spectra for Urine Analysis. Anal Chem 91, 12021–12029 (2019).

32. Xing, S., Shen, S., Xu, B., Li, X. & Huan, T. BUDDY: molecular formula discovery via bottom-up MS/MS interrogation. Nature Methods 2023 20:6 20, 881–890 (2023).

33. Vinaixa, M. et al. Mass spectral databases for LC/MS- and GC/MS-based metabolomics: State of the field and future prospects. TrAC Trends in Analytical Chemistry 78, 23–35 (2016).

34. de Jonge, N. F. et al. MS2Query: reliable and scalable MS2 mass spectra-based analogue search. Nat Commun 14, 1752 (2023).

35. Frainay, C. et al. Mind the Gap: Mapping Mass Spectral Databases in Genome-Scale Metabolic Networks Reveals Poorly Covered Areas. Metabolites 2018, Vol. 8, Page 51 8, 51 (2018).

36. Hoang, C. et al. Tandem Mass Spectrometry across Platforms. Anal Chem 96, 5478–5488 (2024).

37. Kind, T. et al. Identification of small molecules using accurate mass MS/MS search. Mass Spectrom Rev 37, 513 (2017).

38. Böcker, S. Searching molecular structure databases using tandem MS data: are we there yet? Curr Opin Chem Biol 36, 1–6 (2017).

39. Dührkop, K., Shen, H., Meusel, M., Rousu, J. & Böcker, S. Searching molecular structure databases with tandem mass spectra using CSI:FingerID. Proceedings of the National Academy of Sciences 112, 12580–12585 (2015).

