## Supporting Information for "ChemEmbed: A deep learning framework for metabolite identification using enhanced MS/MS data and multidimensional molecular embeddings"

**Supplementary Table 1**. Summary of MS/MS spectra distribution across different spectral databases in positive ionization mode ($[M+H]^+$ adduct), detailing the total number of individual spectra and the corresponding number of compounds.

| Database | Individual spectra | Number of compounds |
|---|---|---|
| NIST20 | 285,550 | 20,818 |
| Agilent Metlin | 37,631 | 9,248 |
| GNPS | 187,005 | 16,102 |
| MSDIAL | 173,478 | 11,870 |

**Supplementary Table 2**. Summary of MS/MS spectra distribution across different spectral databases in positive ionization mode ($[M-H]^-$ adduct), detailing the total number of individual spectra and the corresponding number of compounds.

| Database | Individual spectra | Number of compounds |
|---|---|---|
| NIST20 | 90,113 | 7,798 |
| Agilent Metlin | 14,030 | 3,212 |
| GNPS | 29,566 | 5,006 |
| MSDIAL | 24,673 | 3,561 |

**Supplementary Table 3**. Summary of individual and merged MS/MS spectra and the corresponding unique compounds in positive ionization mode ($[M+H]^+$ adduct), with database prioritization in the order: NIST20 > Agilent METLIN > GNPS > MSDIAL.

| Database | Individual spectra | Merged spectra | Unique compounds |
|---|---|---|---|
| NIST20 | 283,993 | 20,744 | 20,744 |
| Agilent Metlin | 37,007 | 5,823 | 5,823 |
| GNPS | 175,475 | 10,710 | 10,710 |
| MSDIAL | 162,395 | 1,195 | 1,195 |

**Supplementary Table 4**. Summary of individual and merged MS/MS spectra and the corresponding unique compounds in negative ionization mode ($[M-H]^-$ adduct), with database prioritization in the order: NIST20 > Agilent METLIN > GNPS > MSDIAL.

| Database | Individual spectra | Merged spectra | Unique compounds |
|---|---|---|---|
| NIST20 | 89,869 | 7,782 | 7,782 |
| Agilent Metlin | 13,556 | 2,083 | 2,083 |
| GNPS | 27,179 | 3,197 | 3,197 |
| MSDIAL | 22,423 | 1,106 | 1,106 |

### Supplementary Table 5.

| Dataset | Accuracy (top 5) |
|---|---|
| Individual spectra | 55.8% |
| Individual spectra + NL | 59.8% |
| NL alone from merged spectra | 69% |

NL=neutral loses

### Supplementary Table 6.

| Encoding method | Accuracy (Top 1; cosine) | Accuracy (Top 5; cosine) |
|---|---|---|
| Mol2vec embedding | 43% | 76% |
| Fingerprint | 39% | 73% |

### Supplementary Table 7

| Dataset | Ionization & Adduct | Nº Molecules | ChemEmbed accuracy (Top 5) | SIRIUS 6 accuracy (top 5) |
|---|---|---|---|---|
| CASMI 2022 | Positive $[M+H]^+$ | 107 | 33.0 | 28.0 |
| CASMI 2022 | Negative [M-H]- | 64 | 30.0 | 17.0 |
| CASMI 2016 | Positive $[M+H]^+$ | 27 | 66.7 | - |
| CASMI 2016 | Negative [M-H]- | 30 | 61.3 | - |

### Supplementary Table 8

| ARUS dataset | Nº spectra | Nº candidates (cos ≥ 0.95 and Euclidian distance ≤ 25) |
|---|---|---|
| Urine (positive) | 61.767 | 16.030 |
| Urine (negative) | 35.756 | 7.853 |
| Plasma (positive) | 19.422 | 4.221 |
| Plasma (negative) | 15.242 | 2.683 |

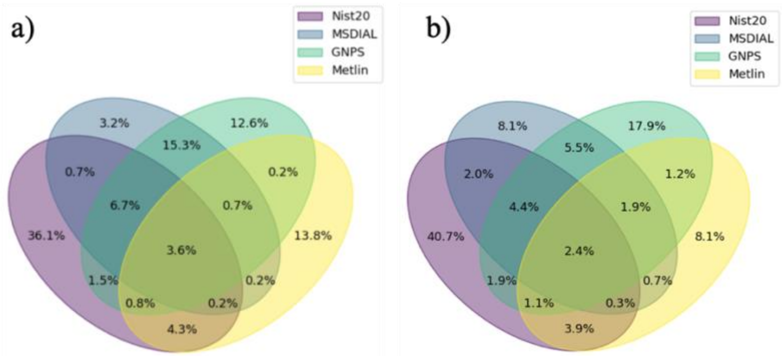


**Supplementary figure 1**. Venn diagrams illustrating the overlap of compounds, identified by their first 14 InChIKey characters, across the four databases. (a) Overlap for positively ionized compounds ($[M+H]^+$). (b) Overlap for negatively ionized compounds ($[M-H]^-$).

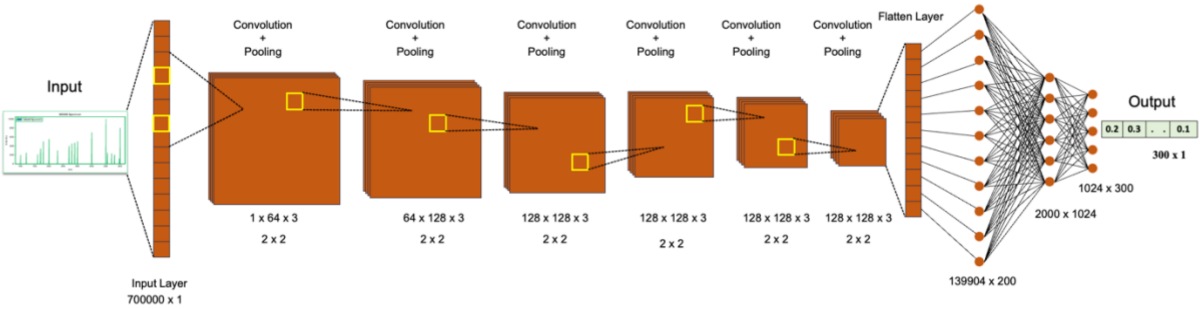


**Supplementary Figure 2.** CNN architecture for processing MS/MS spectra into 300-dimensional feature embeddings. The network consists of six convolutional layers to capture spatial relationships among m/z values, followed by six max pooling layers for dimensionality reduction. A fully connected layer flattens the spatial features, and a final fully connected layer outputs the 300-dimensional Mol2vec embedding. The input is a 70,000-bin vector representing m/z values across the spectrum. Training was conducted with a mini-batch size of 32, using mean squared error as the cost function: 70 epochs for the merged spectra network and 15 epochs for the individual spectra network.

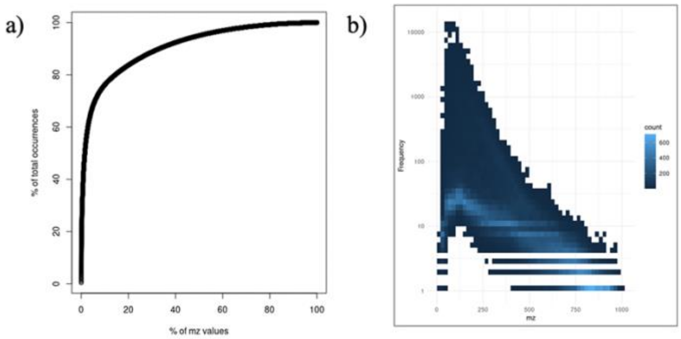


**Supplementary Figure 3**. (a) Skewed distribution of m/z values ranked by their frequency in the MS/MS spectra. The cumulative frequency sum plotted against bin rank percentiles reveals an 80/20 Pareto distribution, where a small fraction of bins accounts for the majority of occurrences. (b) Two-dimensional distribution of bin frequency and m/z value. The plot highlights the prevalence of highly frequent, low-m/z bins that dominate the fragment composition of MS/MS spectra. In contrast, bins with m/z values above 700 Da are much less frequent, with most appearing fewer than 10 times.
